# CDPK2A and CDPK1 form a signaling module upstream of *Toxoplasma* motility

**DOI:** 10.1101/2022.07.19.500742

**Authors:** Emily Shortt, Caroline G. Hackett, Rachel V. Stadler, Gary E. Ward, Sebastian Lourido

## Abstract

The transition between parasite replication and dissemination is regulated in apicomplexan parasites by fluctuations in cytosolic calcium concentrations, effectuated by calcium-dependent protein kinases (CDPKs). We examined the role of CDPK2A in the lytic cycle of *Toxoplasma*, analyzing its role in the regulation of cellular processes associated with parasite motility. We used chemical-genetic approaches and conditional depletion to determine that CDPK2A contributes to the initiation of parasite motility through microneme discharge. We demonstrate that the N-terminal extension of CDPK2A is necessary for the protein’s function. Conditional depletion revealed an epistatic interaction between CDPK2A and CDPK1, suggesting that the two kinases work together to mediate motility in response to certain stimuli. This signaling module appears distinct from that of CDPK3 and PKG, which also controls egress. CDPK2A is revealed as an important regulator of the *Toxoplasma* kinetic phase, linked to other kinases that govern this critical transition. Our work uncovers extensive interconnectedness between the signaling pathways that govern parasite motility.

**IMPORTANCE:** This work uncovers interactions between various signaling pathways that govern *Toxoplasma gondii* egress. Specifically, we compare the function of three canonical calcium dependent protein kinases (CDPKs) using chemical-genetic and conditional-depletion approaches. We describe the function of a previously uncharacterized CDPK, CDPK2A, in the *Toxoplasma* lytic cycle, demonstrating it contributes to parasite fitness through regulation of microneme discharge, gliding motility, and egress from infected host cells. Comparison of analog-sensitive (AS) kinase alleles and conditionally-depleted alleles uncovered epistasis between CDPK2A and CDPK1 implying a partial functional redundancy. Understanding the topology of signaling pathways underlying key events in the parasite life cycle can aid in efforts targeting parasite kinases for anti-parasitic therapies.

## INTRODUCTION

Apicomplexan pathogens like *Toxoplasma gondii, Plasmodium* spp., and *Cryptosporidium* spp. sense and adapt to changes in the host environment throughout their life cycles. These transitions between states often require rapid cellular responses, for which calcium ions (Ca^2+^) are well suited as second messengers. Cells maintain low Ca^2+^ concentrations in the cytosol, in stark contrast to the extracellular milieu and the lumen of some organelles (1). Increased membrane permeability can therefore quickly change the cytosolic Ca^2+^ concentration and initiate signaling. In apicomplexans, such signaling controls progression through the lytic cycle.

The *Toxoplasma* lytic cycle comprises two main phases: a replicative phase during which parasites divide in a parasitophorous vacuole and a kinetic phase that includes egress from the infected host cell, gliding motility, and active invasion of a new host cell. Ca^2+^ signaling mediates the transition between the replicative and kinetic phases of the lytic cycle. After several rounds of replication, parasites actively disrupt surrounding membranes and move out of the infected cell in a process termed egress. This process can be artificially triggered through the use of Ca^2+^ ionophores like ionomycin and A23187 (2–5). Ca^2+^ fluxes can be observed with fluorescent dyes and genetically encoded reporters, which show Ca^2+^ surges that precede gliding motility and invasion in *Toxoplasma (5–7)*. Ca^2+^ oscillations also occur throughout the *Plasmodium* lytic cycle, with peaks in Ca^2+^ concentration preceding microneme secretion, invasion, gliding, and egress (8–11). Control of cytosolic Ca^2+^ concentrations is therefore critical for the regulation of the parasite lytic cycle.

Cytosolic Ca^2+^ surges originate from the release of intracellular stores or by crossing the plasma membrane (PM) from the extracellular space (12, 13). In *T. gondii*, intracellular stores include the endoplasmic reticulum (ER), acidocalcisomes, and plant-like vacuole—with the ER or a related compartment representing the most likely sources of Ca^2+^ during signaling (14). The channels responsible for Ca^2+^ release have yet to be identified in apicomplexans; however, stimulation of the cGMP signaling pathway triggers this process in *T. gondii* and *Plasmodium* spp. (7, 15–17). Treatment with the phosphodiesterase inhibitors zaprinast and BIPPO can block hydrolysis of cGMP and trigger egress (18–20) through the release of intracellular Ca^2+^ stores (7, 15–17). Protein Kinase G (PKG) is a key mediator of Ca^2+^ release, presumably through activation of phosphoinositide signaling, whereby phosphatidylinositol 4,5-bisphosphate (PIP2) is hydrolyzed by phosphoinositide phospholipase C (PI-PLC) into inositol triphosphate (IP3) and diacylglycerol (DAG). This represents a potential branch point in the signaling pathway, with IP3 triggering release of intracellular Ca^2+^ stores through an undefined channel, while DAG is converted into phosphatidic acid (PA) and independently contributes to the kinetic phase (21). Parasite motility therefore requires active cGMP and Ca^2+^ pathways, consistent with the observation that stimulation by Ca^2+^ ionophores cannot overcome the inhibition of PKG in the context of egress (19).

Many Ca^2+^-mediated phenotypes can be attributed to the Ca^2+^-dependent discharge of micronemes, which are specialized organelles that contain adhesins necessary for gliding motility (22, 23). While synthetic treatments like ionophores or phosphodiesterase inhibitors can artificially raise cytosolic Ca^2+^ concentrations and trigger microneme discharge, studies suggest that serum albumin may be a natural agonist of this process (16). The single-pass transmembrane protein MIC2 is a prototypical adhesin that has been used to monitor *T. gondii* microneme discharge. Following its release, MIC2 can be engaged by the actomyosin machinery for gliding motility (12, 24, 25). Several proteases rapidly shed the ectodomain of MIC2 from the surface of the parasite, so its presence in supernatants can be used as a measure of microneme discharge (24). The *P. falciparum* ortholog of MIC2, TRAP, is required for sporozoite motility and invasion (26), and several other adhesins are similarly released and processed in both *Toxoplasma* and *Plasmodium (24, 27, 28). T. gondii* micronemes additionally contain the pore-forming protein PLP1, which permeabilizes the parasitophorous vacuole membrane (PVM) and host PM during egress (29). Following membrane disruption, parasites employ gliding motility to ultimately escape from the ruptured vacuole (25, 30). Perforin-like proteins are also required for egress of *P. falciparum* and *P. berghei* merozoites and gametocytes (31–33). Although the repertoire of microneme proteins differs between species, the Ca^2+^-dependent discharge and participation in gliding motility appear conserved across the phylum (34–36).

Ca^2+^ regulates several cellular processes besides microneme discharge, as evidenced by the wide array of proteins that respond directly to Ca^2+^ concentrations through changes in conformation, stability, localization, or interactions (14). Specialized domains, such as EF hands and C2 domains, endow proteins with the ability to alter their conformation in response to Ca^2+^ binding. Apicomplexans encode several EF hand–containing proteins including calmodulins (CaMs), calcineurin B, and calcium-dependent protein kinases (CDPKs). CDPKs are critical components of the Ca^2+^ signaling network due to their ability to directly respond to Ca^2+^ and phosphorylate other proteins. Although initially identified in plants, CDPKs were later found in ciliates and apicomplexans (14). Despite their similarity to Ca^2+^/CaM–dependent protein kinases, CDPKs are absent from mammals (37). Canonical CDPKs have four C-terminal calmodulin-like EF hands acting as the calcium-binding domain, linked by an autoinhibitory domain to the kinase domain (37, 38). CDPKs in plants control responses to a broad array of stresses, extent of starch accumulation, cell morphology, and viability. Plant CDPKs are expressed across a variety of tissue types and with various subcellular localizations (38). CDPKs exhibit a wide range of affinities for Ca^2+^. For example, soybean CDPKα is activated by ten times lower Ca^2+^ concentrations than CDPKγ (39, 40). Different CDPKs may therefore be tuned to respond to Ca^2+^ spikes of varying magnitudes leading to variable downstream effects that may be further refined by subcellular localization. The array of apicomplexan CDPKs may analogously contribute to the magnitude and compartmentalization of Ca^2+^ responses.

CDPKs are overrepresented in apicomplexan genomes. There are six canonical CDPKs in *T. gondii*, which can be further categorized by having short or long N-terminal extensions (37). Myristoylation sites cap the short extensions of CDPK1 and CDPK3, with an additional palmitoylation site localizing CDPK3 to the parasite PM (19, 41). By contrast, the purpose of the N-terminal extensions remains unknown. A further nine CDPKs in *T. gondii* display non-canonical configurations, with varying numbers and arrangements of EF hands and additional domains (37, 42). The function of most non-canonical CDPKs remains obscure in *T. gondii (42)*, with the exception of CDPK7, which has been implicated in parasite cell division and is critical for phospholipid synthesis and vesicular trafficking (43, 44). By contrast, several canonical CDPKs are required for specific life cycle stages in apicomplexans. In *Plasmodium* spp., several CDPKs regulate specific steps of sexual differentiation (45, 46). CDPKs are also important for kinetic-phase phenotypes in *Toxoplasma* and *Plasmodium*, including microneme discharge, gliding motility, invasion, and egress. TgCDPK1 and TgCDPK3 control *Toxoplasma* egress downstream of PKG (19, 41, 47), analogously to PfCDPK5 in *Plasmodium (10, 48)*. In *Toxoplasma*, CDPK1 is required for invasion as well as egress (19, 47), potentially due to its regulation of microneme secretion under a broader array of conditions than CDPK3. Altogether, CDPKs have been revealed as essential components of the Ca^2+^ signaling network, although the functions of many individual kinases remain unexplored.

In reviewing the results of a genome-wide essentiality screen, most canonical CDPKs were dispensable in tachyzoites; however, CDPK2A remained uncharacterized despite its impact on parasite fitness (49). We sought to characterize the function of CDPK2A within the lytic cycle, comparing its role to that of CDPK1 and CDPK3, two previously-described regulators of the kinetic phase. By turning to a combination of chemical and genetic manipulations we uncover significant interconnectedness between the relevant pathways. Our efforts start to discern the contribution of the various components to the cellular pathways that control microneme discharge, motility, and egress.

## RESULTS

### Chemical-genetic analysis of CDPK2A demonstrates it is involved in parasite egress

To identify the CDPKs that are necessary during the parasite lytic cycle, we examined the results of a genome-wide knockout screen that measured the relative contribution of each gene to fitness as parasites replicated in human fibroblasts (49). Most canonical CDPKs were dispensable in this analysis, including the previously-studied CDPK3; however, CDPK1 and CDPK2A were fitness-conferring, indicated by phenotype scores of -3.3 and -2.05, respectively (**Fig. 1A**). Of these potentially-essential CDPKs, the function of CDPK2A has not been previously examined. Studies of CDPK1 and CDPK3 used analog-sensitive kinase alleles, in which the gatekeeper residue of the ATP-binding pocket of the kinase of interest is mutated to Gly, yielding a binding pocket that accommodates bulky ATP-analog inhibitors (also known as bumped kinase inhibitors), such as 3-MB-PP1 (**Fig. 1B**) (50–52). CDPK1 has a naturally-occurring Gly gatekeeper and can be rendered resistant to 3-MB-PP1 inhibition through mutation of the gatekeeper residue to Met—the same residue that renders CDPK3 and CDPK2A naturally resistant to the inhibitor (50, 51). Comparison of analog-sensitive and insensitive alleles can be used to isolate the effect of inhibiting the kinase in question. Using this approach, CDPK1 was shown to control parasites’ ability to perform gliding motility, invade, and egress from host cells (47). By contrast, compound-mediated inhibition of CDPK3 was used to determine its contribution to motility and egress in response to specific agonists, but not invasion (19), corroborating genetic studies (41, 53).

**Figure 1.**
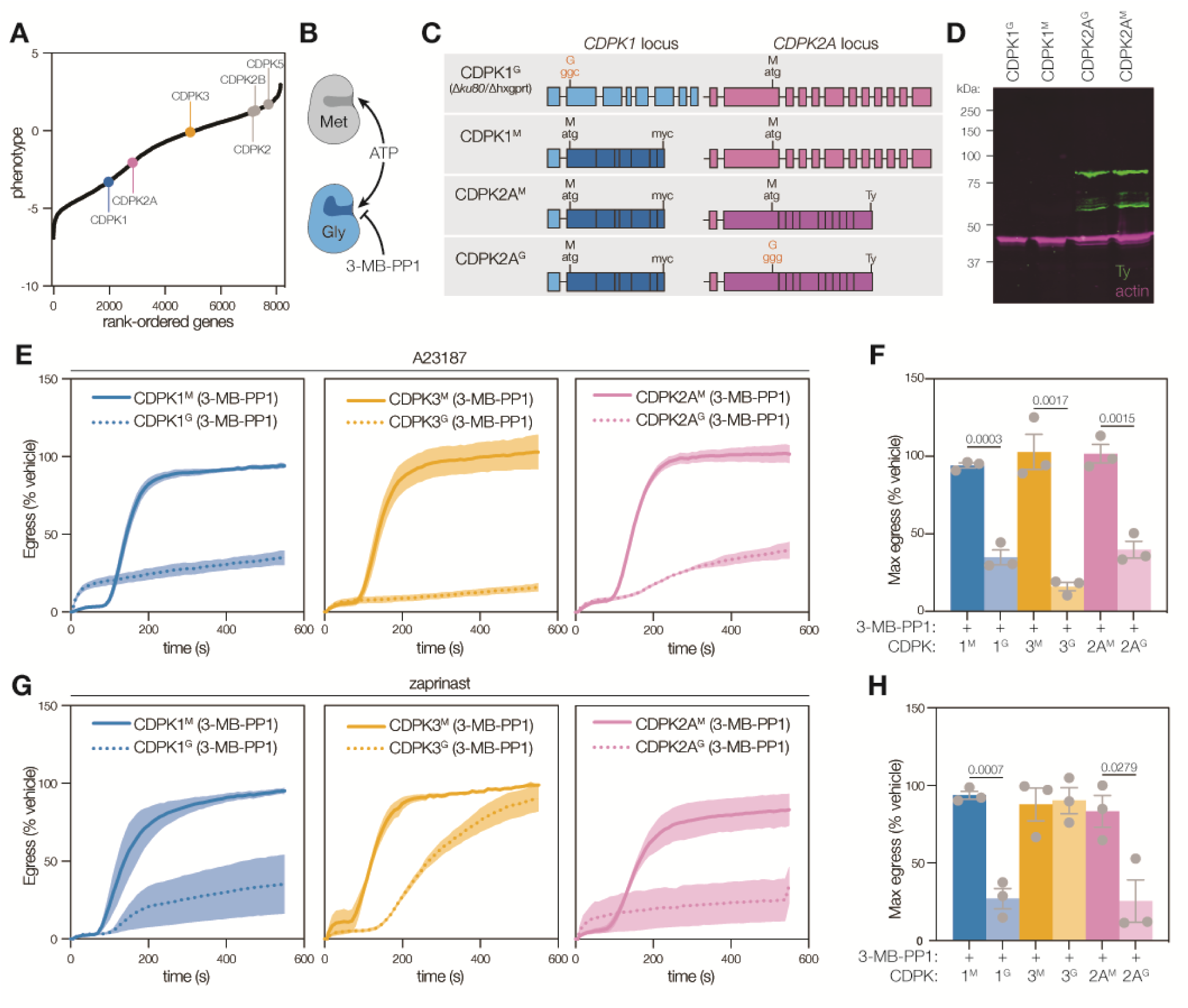
CDPK2A is required for stimulated parasite egress in addition to CDPK1 and CDPK3. **A**. *Toxoplasma* genes ranked by their phenotype, as determined in a genome-wide knockout screen (49). Points represent the mean phenotype score of *n* = 4 replicates. Six canonical calcium-dependent protein kinases are highlighted: CDPK1 (blue), CDPK3 (yellow), and CDPK2A (pink). **B**. Schematic representation of chemical-genetics strategy. The gatekeeper residue of a kinase of interest is mutated, with minimal impact to ATP binding but altered binding of bulky ATP-mimetic inhibitors (e.g., 3-MB-PP1) depending on the size of the binding pocket. **C**. Schematic representation of CDPK1 and CDPK2A loci in the analog-sensitive (AS) kinase strains. Darker shading indicates synthetic DNA. 3-MB-PP1–sensitive alleles are highlighted in red. **D**. Immunoblot of wildtype (CDPK1^G^), parental (CDPK1^M^), and Ty-tagged CDPK2A AS-kinase strains shows expression of the tagged kinase; ACT1 is used as a loading control. **E**. Kinetic traces of A23187-stimulated parasite egress. Graphs are egress of 3-MB-PP1–treated parasites as % of vehicle. A23187 was added 1 s after the start of imaging. Line plots represent the mean ± SEM for *n* = 3 biological replicates. **F**. Maximum egress achieved by each strain during the 10-minute observation window, displayed as % of vehicle. Bars represent the mean ± SEM of *n* = 3 biological replicates; significance calculated by unpaired *t*-test. **G**. Kinetic traces of zaprinast-stimulated parasite egress. Graphs are egress of 3-MB-PP1–treated parasites as % of vehicle. Zaprinast was added 1 s after the start of imaging. Line plots the mean ± SEM for *n* = 3 biological replicates. **H**. Maximum egress achieved by each strain during the 10-minute observation window, displayed as % of vehicle. Bars represent the mean ± SEM of *n* = 3 biological replicates; significance calculated by unpaired *t*-test.

Given the apparent fitness contribution of CDPK2A, we sought to examine its role in the *Toxoplasma* lytic cycle through the use of analog-sensitive alleles. In a strain harboring a 3-MB-PP1–resistant, myc-tagged CDPK1 allele (CDPK1^M^) (19), we modified the CDPK2A locus to introduce a C-terminal Ty epitope tag and either a Met or Gly gatekeeper residue (**Fig. 1C**). The Met modification (CDPK2A^M^) retains the natural resistance of CDPK2A to 3-MB-PP1, whereas the Gly modification (CDPK2A^G^) renders the kinase 3-MB-PP1-sensitive. We confirmed the presence of the mutated gatekeeper residues by allele-specific PCR (**Fig. S1A**) and the expression of the Ty-tagged CDPK2A alleles by immunoblot (**Fig. 1D**). Construction of the isogenic strains carrying alleles of CDPK2A with different susceptibilities to bumped kinase inhibitors allowed us to examine the effect of kinase inhibition on various steps within the lytic cycle.

Parasites can be stimulated to egress from host cells by treating cultures with the calcium ionophore A23187(2) or phosphodiesterase inhibitors such as zaprinast (19) or BIPPO (18). Different agonists have been used to identify pathway-specific requirements for certain kinases (19, 54). Egress leads to loss of host-cell integrity, which can be assayed quantitatively and kinetically as the incorporation of the fluorescent dye 4⍰,6-diamidino-2-phenylindole (DAPI) into the nuclei of permeabilized cells (55). We compared the impact of inhibiting CDPK1, CDPK3, or CDPK2A on A23187-stimulated egress. All three kinases appeared to be required for egress under these conditions. Inhibition of either CDPK1^G^ or CDPK3^G^ strains reduced egress by 66% and 84% compared to vehicle treatment, in agreement with previous findings (19, 41, 47, 53). Analogously, inhibition of CDPK2A^G^ decreased egress by 61% compared to vehicle treatment (**Fig. 1E–F**). Importantly, parasites expressing the resistant allele (CDPK2A^M^) egressed normally in the presence of 3-MB-PP1. To compare the rates of egress, we determined the time required for each strain to achieve half of the maximum egress observed for the respective vehicle-treated control (T_half-max_). Inhibition of all three kinases prevented parasites from reaching half of the maximum egress during the 10-minute observation period (denoted as T_half-max_ > 600 s; **Fig. S1B**). To ensure that the effects observed were the result of differences in egress and not multiplicity of infection, we verified that monolayers had equivalent numbers of parasite vacuoles (**Fig. S1C**). Taken together, these results suggest that CDPK2A contributes to egress, along with CDPK1 and CDPK3.

We next assessed the contribution of CDPK2A to parasite egress after treatment with the phosphodiesterase inhibitor zaprinast. Inhibition of CDPK2A or CDPK1 caused a significant reduction in egress (**Fig. 1G–H**). By contrast, CDPK3 inhibition simply delayed egress (T_half-max_ of 257 s versus 117 s for vehicle-treated parasites), with parasites eventually reaching levels of egress equivalent to CDPK3^M^ (**Fig. 1G–H, Fig. S1D**). These differences were attributable to egress because similar numbers of vacuoles were present in all samples (**Fig. S1E**). The findings are consistent with previous work suggesting that PKG activation, through inhibition of the cGMP-degrading phosphodiesterases, can compensate for inhibition of CDPK3 (19, 54).

### The N-terminal extension of CDPK2A impacts its localization and function

CDPK2A has a long N-terminal extension, which is absent from CDPK1 and CDPK3 (37). We examined whether the N-terminal extension is required for CDPK2A function. Attempts to amplify the 5 end of *CDPK2A* from cDNA yielded two isoforms, which were used to clone two complementation constructs expressed under the heterologous *SAG1* promoter. Complement 1 (c.1) and complement 2 (c.2) differ in how much of the N-terminal extension they include due to alternative splicing; however, both constructs contain the entire kinase domain (**Fig. 2A**). We also generated complement 3 (c.3), which included 1.5 kb of sequence upstream of the predicted translational start site, as well as the first intron, enabling both isoforms to be expressed under endogenous regulation. Complementing alleles had Met gatekeepers and C-terminal HA tags and were expressed in trans in CDPK2A^G^ parasites (**Fig. 2A, Fig. S2A**). This strategy enables inhibition of the endogenous CDPK2A^G^ to assess the functionality of the second copy. The selected clones ---exhibited comparable levels of CDPK2A^M^ expression (**Fig. 2B**). We found that endogenously-tagged CDPK2A (Ty) localizes to the periphery of intracellular parasites, possibly to the inner membrane complex or PM (**Fig. 2C**). Out of the three complementing vectors, only c.3, which retained the endogenous 5⍰ end of the *CDPK2A* mRNA, localized to the parasite periphery, similarly to the endogenous copy. By contrast, c.1 and c.2 localized to the parasite cytosol (**Fig. 2C**). This suggested that localization to the parasite periphery is dependent on the endogenous 5 end of the gene, though not necessarily the presence of the N-terminal extension, perhaps due to a cryptic alternative start site or the precise timing of expression.

**Figure 2.**
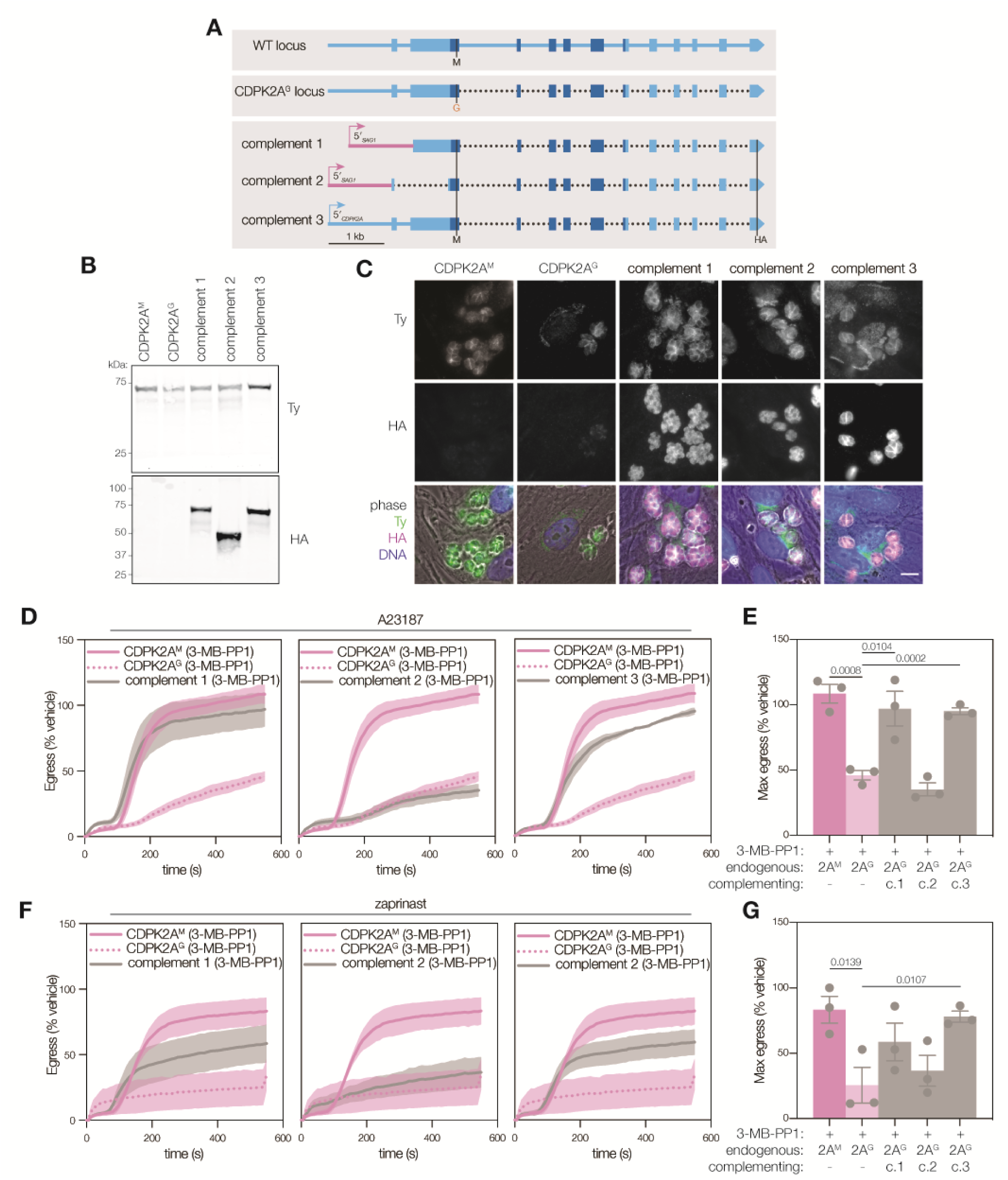
The N-terminal extension of CDPK2A influences its localization and is necessary for parasite egress. **A**. Schematic representation of complementing constructs with comparison to CDPK2A genomic locus. Dotted lines indicate gaps in sequence where introns have been removed. Shaded region encoding the kinase domain. **B**. Immunoblot of complemented strains probing for the endogenous allele with Ty and the complementing allele with HA. **C**. Immunofluorescence microscopy of endogenously Ty-tagged CDPK2A^M^ and CDPK2A^G^ and HA-tagged complementing copies of CDPK2A^M^. Merged image displays Ty (green), HA (magenta), DNA (blue), and phase (greyscale). Scale bar is 10 μm. **D**. Kinetic traces of A23187-stimulated parasite egress. Graphs are egress of 3-MB-PP1–treated parasites as % of vehicle. A23187 was added 1 s after the start of imaging. Line plots represent mean ± SEM for *n* = 3 biological replicates. **E**. Maximum egress achieved by each strain during the observation window, displayed as % of vehicle. Bars represent mean ± SEM of *n* = 3 biological replicates; significance calculated by unpaired one-tailed *t*-test. **F**. Kinetic traces of zaprinast-stimulated parasite egress. Graphs are egress of 3-MB-PP1–treated parasites as % of vehicle. Zaprinast was added 1 s after the start of imaging. Line plots represent mean ± SEM for *n* = 3 biological replicates. **G**. Maximum egress achieved by each strain during the observation window, displayed as % of vehicle. Bars represent the mean ± SEM of *n* = 3 biological replicates; significance calculated by unpaired one-tailed *t*-test.

We next assessed the function of the complementing alleles. We inhibited the endogenous CDPK2A^G^ allele and assessed whether each of the complemented strains could egress following A23187 or zaprinast induction. With A23187 stimulation, c.1 and c.3 strains egressed normally despite inhibition of the endogenous allele; however, inhibitor-treated c.2 parasites exhibited no complementation (**Fig. 2D–E**). The two alleles that did complement (c.1 and c.3) had egress kinetics comparable to 3-MB-PP1–resistant CDPK2A^M^ parasites, reaching half the maximal egress around 200 s (**Fig. S2B**). We confirmed that in all cases the monolayers were equivalently infected (**Fig. S2C**). Complementation during zaprinast-induced egress was intermediate but followed similar trends, with nearly-wild-type responses for c.3, an intermediate response for c.1, and no apparent complementation by c.2 (**Fig. 2F–G**). This partial rescue was also reflected in the rates of egress (**Fig. S2D–E**). These findings suggest that the N-terminal extension of CDPK2A—present in c.1 and c.3 constructs, but absent in c.2—is required for parasite egress. The functional complementation contrasts with the localization of the proteins encoded by these alleles, where only c.1 matched the peripheral localization of protein encoded by the endogenous allele.

### Conditional depletion of CDPK2A only partially mimics chemical inhibition

Off-target effects of 3-MB-PP1 interfere with the chemical-genetic approach described above during long-term culture, making it challenging to assess the impact of kinase inhibition over several lytic cycles (56). Auxin-inducible degradation of target proteins has been adapted to *Toxoplasma* and used to investigate protein kinases and associated signaling pathways (57, 58). We employed this conditional-depletion system as an orthogonal strategy to assess CDPK function. Briefly, a protein of interest is tagged with an auxin-inducible degron (AID) in a strain expressing TIR1. When transgenic parasites are treated with the plant hormone auxin—most commonly 3-indoleacetic acid (IAA)—the tagged protein is ubiquitinated and targeted by the proteasome for degradation (59) (**Fig. 3A**). Protein depletion occurs within minutes to hours in *Toxoplasma*, depending on the protein of interest (57, 58). We generated a panel of strains in which CDPK1, CDPK3, or CDPK2A were tagged at their C termini with the Ty epitope followed by mNeonGreen and a minimal auxin-inducible degron (mAID; **Fig. S3A**). Localization of CDPK2A-AID was consistent with the chemical-genetic alleles, with mNeonGreen signal observed at the parasite periphery (**Fig. 3B**). Expression of all mNeonGreen-tagged alleles was measured by FACS in parasites with and without IAA treatment. Three hours of IAA treatment was sufficient to observe robust and uniform depletion of each CDPK (**Fig. 3C**). Based on these results, we treated the parasites with auxin for a minimum of 3 h for downstream analyses, achieving kinase degradation within less than a single cell cycle.

**Figure 3.**
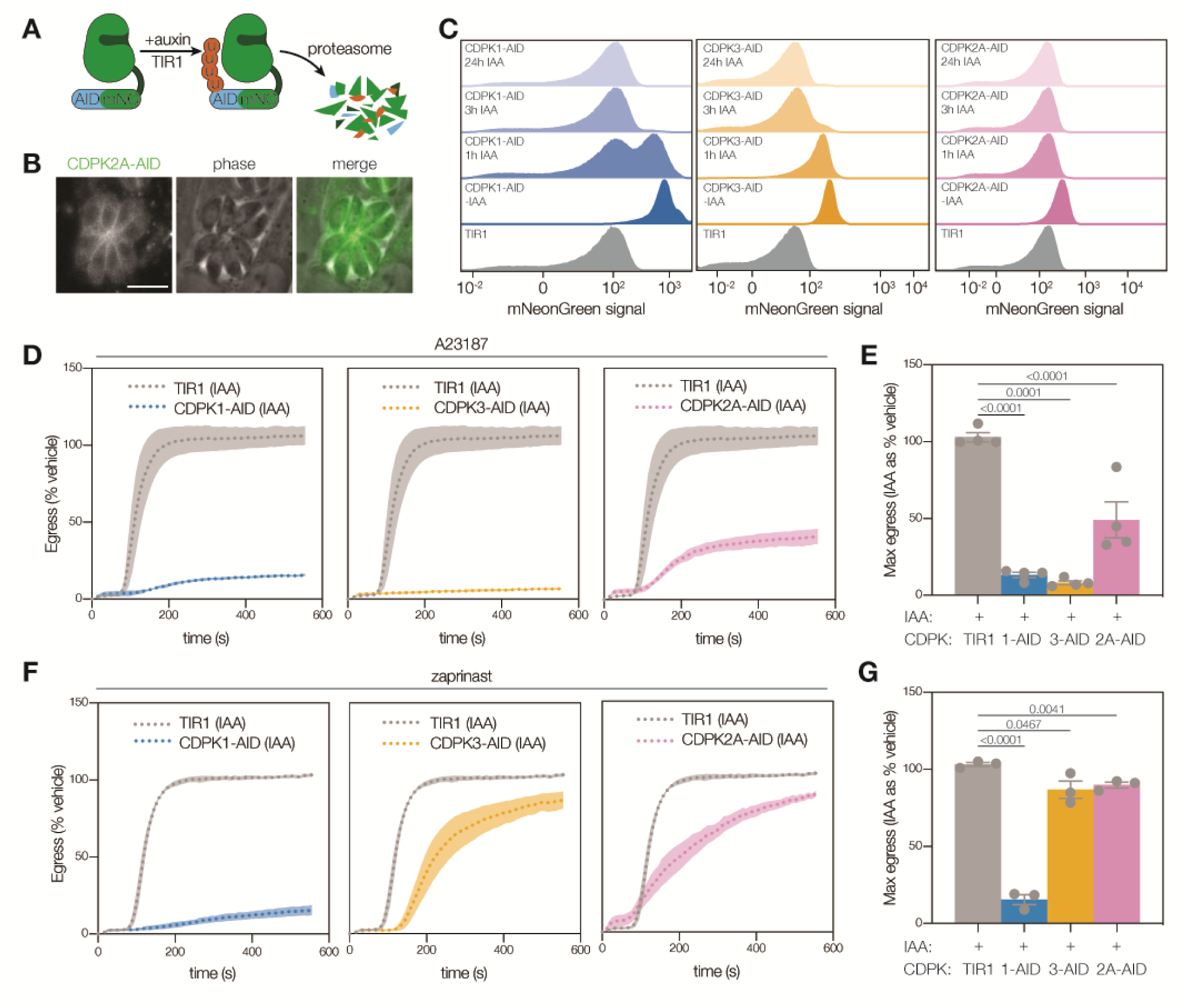
Conditional depletion of CDPKs reveals CDPK2A is required for A23187-stimulated egress, but not for zaprinast-stimulated egress. **A**. Schematic representation of a CDPK tagged with a minimal auxin-inducible degron (mAID), Ty, and mNeonGreen. When IAA is added, the TIR1 ubiquitin ligase complex is recruited to the tagged protein and targets it for proteasomal degradation. **B**. Live-cell imaging of CDPK2A-AID parasites shows the mNeonGreen signal concentrated at the parasite periphery. Scale bar is 10 μm. **C**. FACS analysis of mNeonGreen-tagged CDPK-AID parasites and TIR1 parental line with and without the addition of auxin (IAA) shows expression of mNeonGreen fusion and kinetics of auxin-induced depletion. **D**. Kinetic traces of A23187-stimulated parasite egress. Graphs are egress of 3-MB-PP1–treated parasites as % of vehicle. A23187 was added 1 s after the start of imaging. Line plots represent mean ± SEM for *n* = 4 biological replicates. **E**. Maximum egress achieved by each strain during the observation window, displayed as % of vehicle. Bars represent mean ± SEM of *n* = 4 biological replicates; significance calculated by unpaired *t*-test. **F**. Kinetic traces of zaprinast-stimulated parasite egress. Graphs are egress of 3-MB-PP1–treated parasites as % of vehicle. Zaprinast was added 1 s after the start of imaging. Line plots the mean for *n* = 3 biological replicates with error bands representing SEM. **G**. Maximum egress achieved by each strain during the observation window, displayed as % of vehicle. Bars represent the mean ± SEM of *n* = 3 biological replicates; significance calculated by unpaired *t*-test.

We assessed egress following acute depletion of CDPK1, CDPK3, or CDPK2A. Stimulation with A23187 induced minimal egress in CDPK-depleted parasites, indicating that all three kinases are necessary for egress under these conditions (**Fig. 3D–E**); this is consistent with our chemical-genetic approach. Analysis of egress kinetics showed that all three depleted lines failed to egress within the observation window, in contrast to the TIR1 parental parasites, which achieved half-maximum egress in 115 s (**Fig. S3B–C**).

We next assessed the ability of CDPK-depleted parasites to egress in response to zaprinast stimulation. Depletion of CDPK1 or CDPK3 phenocopied their chemical-genetic inhibition. CDPK1-depleted parasites were unable to egress, whereas CDPK3-depleted parasites displayed delayed but near-complete egress (**Fig. 3F–G**), achieving half-maximum egress at 260 s compared to 120 s for the parental TIR1 strain (**Fig. S3D**). Surprisingly, CDPK2A-depleted parasites achieved normal levels of egress (90% of vehicle-treated parasites), albeit with a delay (T_half-max_ of 230 s), despite equivalent levels of overall infection (**Fig. 3F–G, Fig S3D–E**). We conclude that differences between the strains used for chemical inhibition or depletion render CDPK2A differentially required for egress. Such differences may result from manipulation of the *CDPK2A* locus or the strain background in which the mutants were generated.

### Inhibition of CDPK1 reveals epistasis with CDPK2A

We considered whether differences in the CDPK1 alleles of the parental lines used—CDPK1^M^ for chemical genetics and CDPK1^G^ for conditional depletion—could lead to the differential requirement for CDPK2A in zaprinast-stimulated egress when assessed by either approach. Enzymatic assays have demonstrated that CDPK1^G^ is more catalytically active than CDPK1^M^, although either allele can support parasite replication (60). Based on the presence of CDPK1^M^ in the analogue-sensitive (AS) kinase lines and the observed overlapping phenotypes of CDPK1 and CDPK2A, we hypothesized that partial loss of CDPK1 activity may lead to an increased reliance on CDPK2A for egress.

We assessed whether CDPK1 and CDPK2A exhibit epistasis by broadly examining the lytic cycle during plaque formation. We partially inhibited CDPK1 with sublethal concentrations of 3-MB-PP1 in the context of CDPK2A expression or depletion, using the CDPK2A-AID strain. CDPK2A-depleted parasites failed to form plaques when CDPK1 was partially inhibited. By contrast, plaquing was not impacted by partial inhibition of CDPK1 in the context of CDPK3 depletion (**Fig. 4A**). This suggests that the requirement for CDPK2A during the lytic cycle depends on the level of CDPK1 activity.

**Figure 4.**
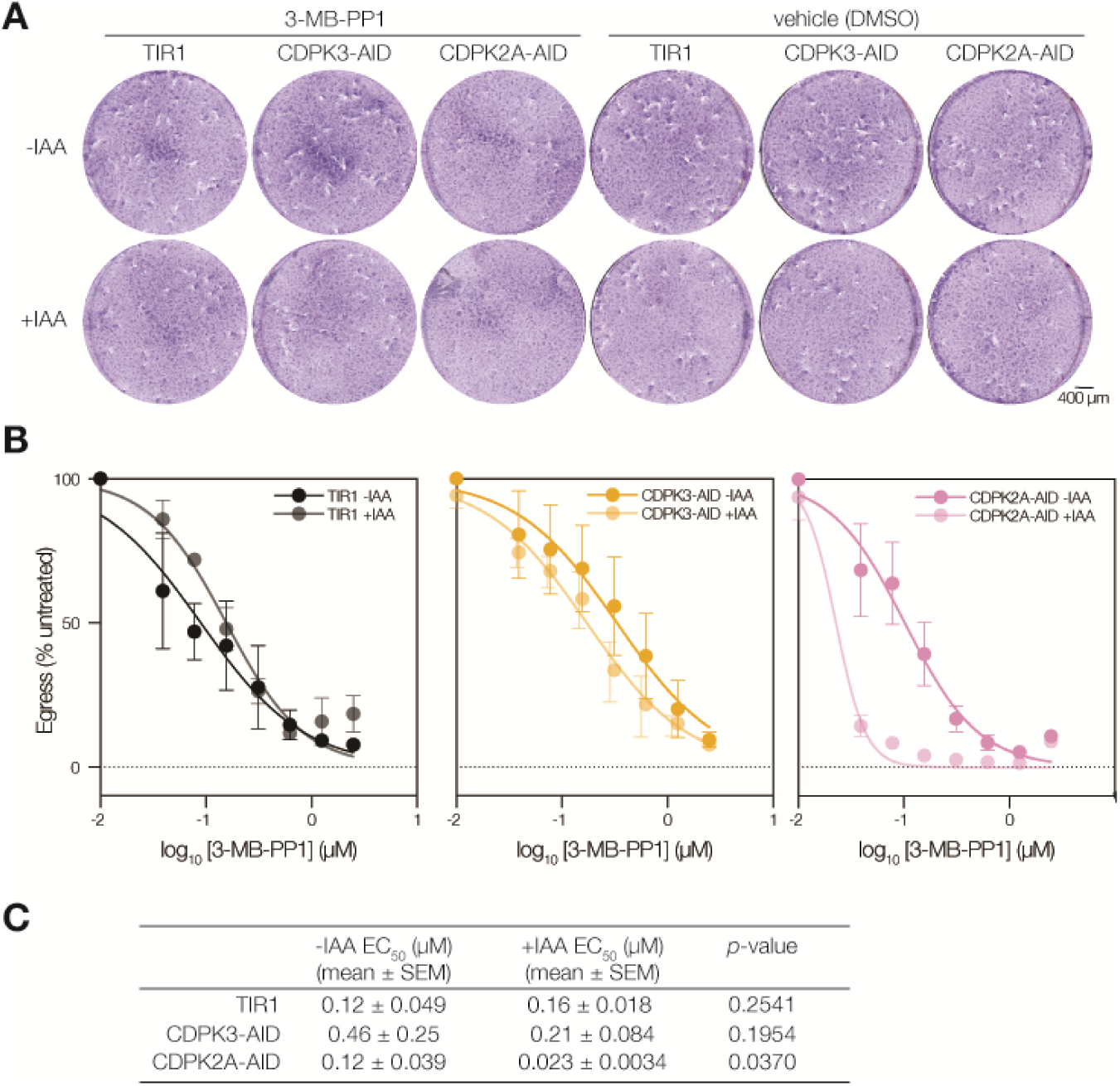
CDPK1 and CDPK2A comprise a signaling module that regulates zaprinast-stimulated egress and the lytic cycle. **A**. Partial inhibition of endogenous CDPK1^G^ by 3-MB-PP1 (40nM) leads to ablation of plaque formation by parasites conditionally depleted of CDPK2A. Scale bar represents 400 μm. **B**. Dose-response of CDPK1 inhibition with 3-MB-PP1 monitoring endpoint zaprinast-stimulated egress in parasites depleted of or expressing the indicated CDPK. Mean ± SEM plotted for *n* = 3 biological replicates. **C**. EC50 (μM) for 3-MB-PP1 of CDPK-depleted or untreated parasites; significance calculated by unpaired one-tailed *t*-test across *n* = 3 biological replicates.

We next examined the epistatic interaction between the two kinases during zaprinast-stimulated egress using endpoint assays. Treating parasites with a range of 3-MB-PP1 concentrations to inhibit CDPK1^G^, we observed that CDPK2A depletion renders parasites hypersensitive to CDPK1 inhibition (**Fig. 4B, Fig. S4**). By contrast, depletion of CDPK3 did not change the sensitivity of parasites to CDPK1 inhibition by 3-MB-PP1; significance was calculated using an F test for fitting all data to a single curve— CDPK2A-AID data points did not fit a single curve (p<0.0001), whereas TIR1 and CDPK3-AID data points, irrespective of IAA treatment, fit to one curve (ns). This observation is supported by the calculated EC50 for 3-MB-PP1, which decreases significantly when CDPK2A-AID parasites are treated with auxin (**Fig. 4C**). The combination of chemical inhibition and protein knockdown allowed us to demonstrate that CDPK2A becomes more critical for egress when CDPK1 activity is compromised due to a mutant gatekeeper allele as in CDPK1^M^ or chemical inhibition. We surmise that the AID-tagged lines more closely reflect physiological conditions—with a wildtype CDPK1 allele—than AS kinase lines, so we examined the function of CDPK2A across the lytic cycle using conditional depletion.

### Conditional depletion reveals a role for CDPK2A at various stages of the lytic cycle

Plaque formation captures repeated cycles of host cell lysis. As expected, depletion of CDPK1 blocked plaque formation (47), whereas CDPK3 appeared completely dispensable (41, 49, 53). Depletion of CDPK2A resulted in the formation of fewer plaques than the vehicle-treated tagged line (**Fig. 5A, Fig. S5A**). These results are in line with the phenotype scores in the genome-wide fitness screen (49), placing CDPK2A at an intermediate fitness contribution, between CDPK1 and CDPK3. Additionally, CDPK2A-AID parasites formed smaller plaques than the parental line, and plaque size further decreased when CDPK2A was depleted by IAA treatment (**Fig. 5B, Fig. S5B**). The observed hypomorphism between CDPK2A-AID and parental TIR1 parasites suggests that AID-tagging of the kinase may impact its function. The impact of CDPK2A depletion on plaquing efficiency may result from impaired or inefficient invasion, slower parasite replication, decreased motility of parasites during successive rounds of lysis, or a combination thereof, which must be deconvoluted through single-phenotype assays.

**Figure 5.**
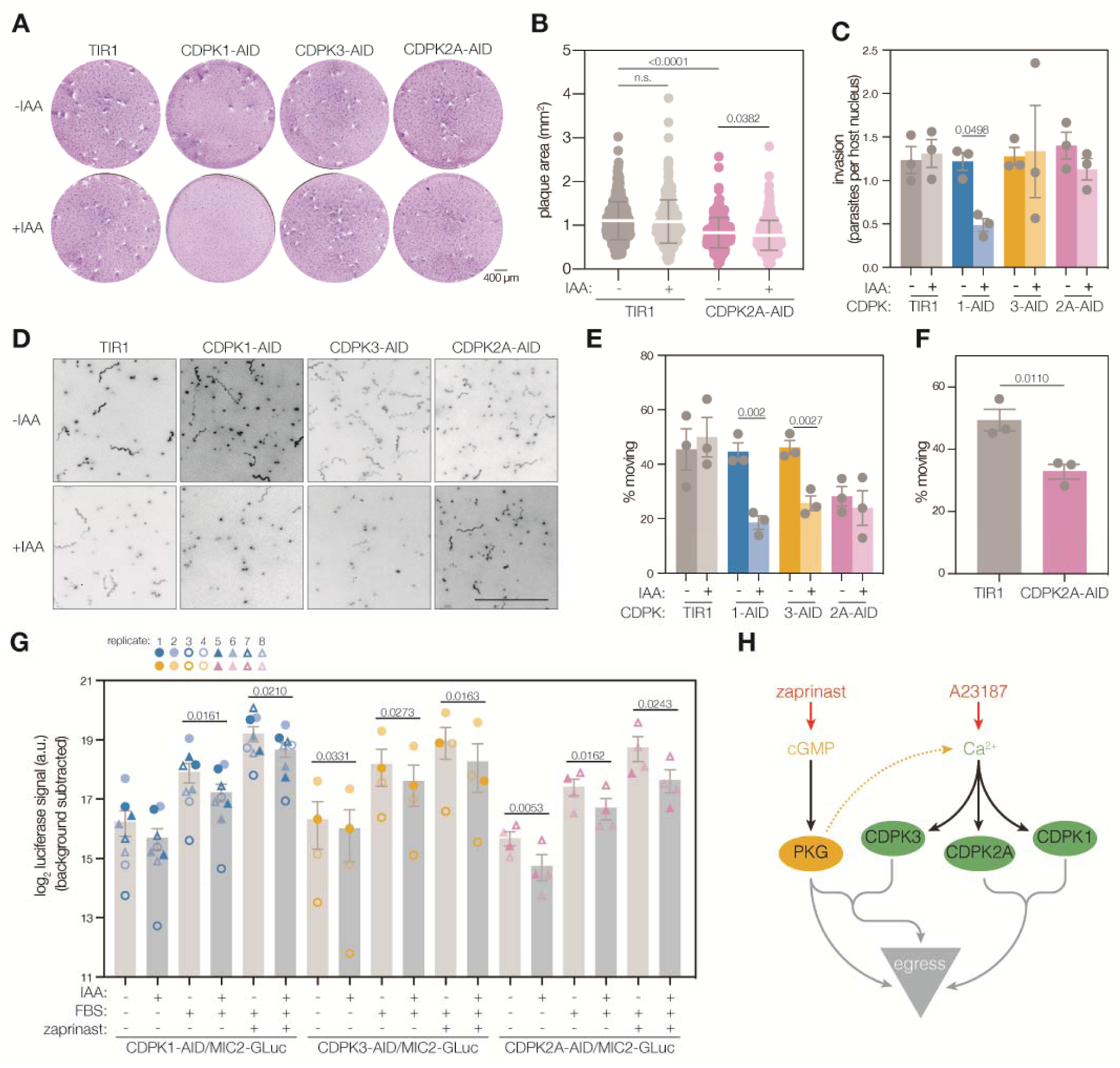
CDPK2A impacts plaque formation, gliding motility, and microneme discharge. **A**. Depletion of each of the three CDPKs impacted plaque formation differently: parasites depleted of CDPK1 formed no plaques, those depleted of CDPK2A formed smaller plaques, and those depleted of CDPK3 plaqued normally. Images are representative of *n* = 4 biological replicates. Scale bar is 400 μm. **B**. CDPK2A-AID parasites form smaller plaques than the TIR1 parental line, and the effect is exacerbated by the depletion of the kinase (+ IAA). Scatter plot displays areas for >270 individual plaques per sample; mean ± SD is overlaid; *p* value calculated by unpaired *t*-test. **C**. CDPK2A depletion does not impact parasites’ ability to invade host cells. Bars represent the mean ± SEM for *n* = 3 biological replicates; *p* value calculated by paired *t*-test. **D**. 3D gliding motility in Matrigel following depletion of each kinase. Maximum intensity projections of tracked parasites. Scale bar is 100 µm. **E**. Proportion of parasites observed moving in 3D gliding motility assays. Each point is the mean of 3 technical replicates with 634-1500 observations per condition; bars represent the mean ± SEM of *n* = 3 biological replicates; significance calculated by unpaired one-tailed *t*-test. **F**. Direct comparison of the proportion of parasites gliding in 3D for TIR1 and CDPK2A-AID parasites without IAA. Each point is the mean of 3 technical replicates with 1011 and 1157 observations for TIR1 and CDPK2A-AID respectively; bars represent the mean ± SEM of *n* = 3 biological replicates; significance calculated by unpaired one-tailed *t*-test. **G**. Depletion of CDPKs decreases microneme discharge. Gaussia-luciferase activity, from a microneme-localized construct, was assayed in supernatants following 30 minutes of stimulation with FBS and zaprinast. Points represent individual biological replicates (*n* = 8 for CDPK1-AID, *n* = 4 for CDPK3-AID or CDPK2A-AID), with bars indicating mean ± SEM; significance calculated by paired one-tailed *t*-test. **H**. Proposed model for the relationship between kinases that positively regulate parasite motility.

We assessed whether CDPK2A is required for parasite invasion of host cells using an immunofluorescence assay that distinguishes between invaded and extracellular parasites. Depletion of CDPK1 blocked parasite invasion; however, depletion of CDPK3 or CDPK2A did not affect parasite invasion (**Fig. 5C**), consistent with previous findings for CDPK1 and CDPK3(19, 41, 47). The dispensability of CDPK2A for invasion was corroborated by chemical-genetics. Inhibition of analog-sensitive alleles of CDPK1 or CDPK2A with 3-MB-PP1 confirmed that CDPK1 is necessary for invasion, whereas CDPK2A is dispensable for this process (**Fig. S5C**).

Gliding motility precedes invasion and is a necessary aspect of parasite egress from host cells, making it critical for parasite spread (61). We quantitatively assessed the 3D motility of CDPK-depleted parasites in Matrigel, which may better capture subtle motility defects that are hard to appreciate by traditional 2D motility assessment (62) (**Fig. 5D**). Depletion of CDPK1 or CDPK3 decreased the percentage of parasites moving during the assay by 58% or 44% respectively. CDPK2A-AID parasites appeared to move less than the parental TIR1 strain, although their motility was not decreased further by pre-treatment with IAA (**Fig. 5E**). We directly compared the motility of TIR1 parental and CDPK2A-AID tagged parasites without the addition of IAA, confirming that CDPK2A-AID tagging decreased parasite motility significantly (**Fig. 5F**). Track length and displacement were significantly altered by addition of IAA to CDPK1-AID parasites only, and maximum and speed mean speed were not altered by IAA treatment in any of the strains analyzed (**Fig. S5D–G**). Consistent with our observations, CDPK1 was previously implicated in parasite motility in two-dimensional analyses (19, 47). There is some ambiguity as to CDPK3’s contribution to parasite motility in previous 2D analyses, depending on whether parasites were stimulated by switching from intracellular to extracellular buffer (19, 53, 63) or by treatment with A23187(41). Nevertheless, it is clear that CDPK3-mediated phosphorylation of MyoA contributes to motility (63). Our results suggest that CDPK2A, along with CDPK1 and CDPK3, plays a critical role in gliding motility; however, AID tagging seems to sufficiently reduce CDPK2A activity such that no further reduction in gliding motility is observed under IAA treatment. We expect that certain phenotypes might require higher levels of kinase activity, revealing the hypomorphism of the conditional allele.

We further assessed the ability of CDPK2A-depleted parasites to secrete micronemal contents. We expressed Gaussia luciferase (GLuc) fused to myc-tagged MIC2 in CDPK-AID transgenic lines in order to study microneme protein secretion following knockdown of each kinase (**Fig. S5H**). Following CDPK depletion, parasites were stimulated to secrete with fetal bovine serum (FBS) alone or supplemented with zaprinast. Excreted/secreted antigen (ESA)-containing supernatants were collected, and MIC2 secretion was measured by luciferase signal. Parasites depleted of CDPK1, CDPK3, or CDPK2A secreted less MIC2 when stimulated with either FBS alone or FBS and zaprinast for 30 min (**Fig. 5G**).

Interestingly, CDPK2A is not required for zaprinast-stimulated microneme protein secretion at an acute 5 min time point, even though CDPK1 and CDPK3 are each required (**Fig. S5I**). We further verified that basal MIC2 levels were equivalent between strains and that CDPK depletion does not impact total MIC2 in parasite lysates (**Fig. S5J**). Overall, these results suggest CDPK1, CDPK3, and CDPK2A are all involved in microneme protein secretion, but to different degrees.

## DISCUSSION

We examined the role of CDPK2A in the lytic cycle of *Toxoplasma*, analyzing its contribution to parasite fitness through processes related to parasitism, including microneme protein secretion, gliding motility, and egress from host cells. Using a combination of chemical-genetic and conditional depletion methods, we show that CDPK2A contributes to the initiation of parasite egress through microneme discharge and gliding motility. We further demonstrated that the N-terminal extension of CDPK2A is necessary for the protein’s function in egress. Contrasting results from chemical inhibition and conditional depletion studies revealed an epistatic interaction between CDPK2A and CDPK1. Our results suggest that CDPK2A and CDPK1 work together to mediate egress following the stimulation of the PKG pathway (**Fig. 5H**). This signaling module might be differentially compartmentalized from CDPK3 and PKG, which localize to the PM. Our work uncovered additional complexity and interconnectedness in the signaling pathways that govern key events during the parasite lytic cycle.

CDPK2A contributes to parasite fitness. Plaque assays showed limited growth for parasites depleted of CDPK2A—an intermediate effect between the dispensable CDPK3 and the essential CDPK1. These observations are consistent with previous genome-wide loss-of-function screens, which had calculated an intermediate phenotype for CDPK2A, between CDPK1 and CDPK3(49). We have sought to determine what differentiates the fitness-conferring CDPK1 and CDPK2A from the dispensable CDPK3. Chemical-genetic approaches previously showed that accumulation of cGMP, which activates PKG, can compensate for loss of CDPK3 during egress (19). The reliance on CDPK2A similarly appeared to be conditional on the activity of other kinases. While all three CDPKs were required for egress in response to calcium ionophores, as with CDPK3, conditional depletion of CDPK2A could be partially compensated through hyperstimulation of the PKG pathway; however, in the case of CDPK2A, compensation depended on the level of CDPK1 activity.

Epistasis appears pervasive among the pathways regulated by CDPKs. As mentioned above, activation of PKG through the application of phosphodiesterase inhibitors (e.g., zaprinast) enables parasite egress despite CDPK3 inhibition or loss (19, 54). Epistasis between PKG and CDPKs has also been observed in *Plasmodium* spp. at various stages of the intraerythrocytic cycle (15, 64, 65). Genetic interaction between PbCDPK4 (the ortholog of TgCDPK1) and PKG was revealed by a *P. berghei* screen (15).

Analogously to our chemical-genetic results, PbCDPK4 becomes critical for parasite invasion and motility in a genetic background expressing a variant of PKG in which the gatekeeper residue has been mutated (PKG^T619Q^) (15). It is inferred from phenotypes associated with PKG activity that the PKG^T619Q^ mutant is less active than wildtype. As with the TgCDPK1^G128M^ allele used in our chemical-genetic approach, these mutants retain sufficient kinase activity to sustain parasite viability, yet the mutation clearly places a strain on other aspects of the signaling network. Such interactions likely extend further into the network, since double knockouts of PbCDPK1 and PbCDPK4 are viable but cannot be generated in parasites expressing PKG^T619Q^ (15). The interconnectivity of CDPK networks may render them more plastic. Indeed, studies in *P. falciparum* suggest that parasites rapidly adapt to the loss of PfCDPK1 activity, perhaps through upregulation of other CDPKs (65, 66).

The plasticity of the signaling networks controlling egress can be manipulated through pharmacological stimuli that obscure or exaggerate the function of individual pathway components, revealing novel connectivity or dependencies. As described above, hyperactivation of the PKG pathway overcomes inhibition of CDPK2A or CDPK3. Analogously, *P. falciparum* parasites deficient in CDPK5 fail to egress, but this block can be overcome by hyperactivation of PKG (10, 48)). The degree of pathway overstimulation, whether through ionophore or phosphodiesterase inhibitor treatment, influences the interpretation of the results—particularly since the natural levels or dynamics of these second messengers are rarely known. Hyperactivation of a pathway may also force interactions that would otherwise not occur at the basal state (67). With this context, we can consider that epistatic interactions may result from shared substrates or the redundancy of independent pathways. PKG has been shown to be a calcium regulator in *T. gondii* and *Plasmodium* spp.(7, 15–17, 68), placing it upstream of CDPK activation. In plants, CDPKs are tuned to respond to different calcium concentrations (38), raising the possibility that dependency on different parasite CDPKs may result from the magnitude of the calcium surge elicited by PKG. Nevertheless, the subcellular sorting of epistatic interactions—with CDPK3 and PKG strictly localized to the PM—argues for potential overlap in their substrates as the mechanism underlying their epistasis.

Our studies also reflect some of the challenges inherent in studying protein kinases and interconnected signaling networks. While the goal is often to infer the role of a kinase in its native state, genetic perturbations may result in compensatory changes that obscure its function. While AID knockdown of CDPK1 and CDPK3 phenocopied chemical-genetic findings for egress, CDPK2A knockdown did not. Surprisingly, the discrepancy between chemical inhibition and AID depletion of CDPK2A could be attributed to the difference in CDPK1 alleles between the two systems, since partial inhibition of CDPK1 results in a stronger requirement for CDPK2A in zaprinast-stimulated egress. This is consistent with biochemical studies that had revealed reduced ATP affinity of TgCDPK1^G128M^ relative to the wildtype enzyme (60). Analogously, phenotypic assays such as 3D gliding motility and plaque size argue for hypomorphism of the CDPK2A-AID allele. Inspection of CDPK2A-AID parasites’ motility tracks suggested they may move along less-tightly-wound (or lower amplitude) corkscrews compared to other strains. An altered geometry of movement may be less efficient, resulting in the smaller plaque areas observed for CDPK2A-AID parasites. Taken together, these results argue for caution in the interpretation of perturbed signaling systems; nevertheless, comparison of multiple approaches can be used to infer the native function of protein kinases like CDPK2A.

Several signaling pathways converge on the regulation of microneme discharge, including those controlled by CDPK2A. Microneme discharge lies upstream of parasite egress, gliding motility, and invasion through the release of diverse proteins, including perforins that disrupt the PVM (29) and adhesins that mediate substrate attachment (69, 70). Depletion of any of the studied CDPKs resulted in a decrease in microneme discharge. The effect of CDPK2A depletion was only evident following prolonged periods of microneme discharge (30 min). By contrast, the effect of CDPK1 and CDPK3 was already evident within 5 min of stimulation. This may suggest a model in which some CDPKs regulate an initial wave of secretion, while others regulate the sustained response. Previous studies of CDPK3-knockout parasites reported normal MIC2 secretion for extracellular parasites stimulated with A23187 or ethanol (41), although intracellular parasites clearly depend on CDPK3 to permeabilize the parasitophorous vacuole upon A23187 treatment (19, 41). Differences in the sensitivity or conditions of our assays (e.g., the use of intracellular buffer in our microneme discharge assays) may have focused our assays on the responses that govern egress. Previous work also reported that the contributions of CDPK1 and CDPK3 to microneme discharge depended on the agonist used (19). Consistent with these observations and the epistatic interactions discussed above, CDPK2A may impact microneme secretion to different extents across the lytic cycle depending on the stimuli experienced by parasites.

We demonstrated that the N-terminal extension of CDPK2A is necessary for its function. Complementing constructs that expressed a full-length version of CDPK2A were able to egress when the endogenous kinase was inhibited. The N terminus may contain localization determinants that drive CDPK2A to the parasite periphery, although the predicted gene model lacks consensus motifs for myristoylation or palmitoylation that participate in the localization of CDPK3 to the PM (19, 41). CDPK2A was also not detected in mass spectrometry datasets enriching for myristoylated (71) or palmitoylated (72) proteins. Proteomic studies have detected CDPK2A peptides spanning the coding sequence of exon 2 (exon 1 in c.1), further suggesting that the truncated c.2 sequence generated from cDNA is not the prominent species in wild-type parasites (71). Only the complementing construct driven by the endogenous 5⍰ UTR and promoter yielded a protein that co-localized with the endogenous CDPK2A, suggesting localization to the parasite periphery depends on endogenous regulatory signals rather than the N terminus of the protein. We cannot exclude that an alternative translation start site is used, giving rise to dually-localized species. Methionine 59 in the longer gene model may be the true start site, matching the predicted coding sequence in *Hammondia hammondi* (73). The existing data suggest that localization to the parasite periphery is not strictly required for CDPK2A’s function, in contrast to CDPK3, which must be peripherally-localized via myristoylation and palmitoylation in order for parasites to egress (19, 41). CDPK orthologs containing N-terminal extensions appear not to universally localize to any given compartment. PfCDPK5, which controls egress, associates with parasite membranes, possibly including the cytosolic-facing side of micronemes (10, 48), while PfCDPK3, required for ookinete motility, is cytoplasmic (11, 74). There is also precedent for signaling-related proteins to express multiple functionally distinct isoforms, often arising from alternative translational initiation. For example, in *E. tenella* and *T. gondii*, one isoform of PKG is N-acylated and localizes to the PM, while the other isoform is cytoplasmic. Interestingly, either isoform can function if targeted to the PM (58, 75). Isoform diversity may further drive the plasticity of CDPK signaling networks, although this has not been formally addressed by our work.

The use of both chemical inhibition and conditional depletion to study CDPK2A function uncovered additional complexity and interconnectedness in the signaling pathways that govern the lytic cycle. We observe that CDPK1, CDPK3, and CDPK2A are all involved to varying degrees in microneme discharge, with functional consequences during egress, gliding motility, and invasion. We also further describe functional redundancy that structures the pathway into two signaling modules that are jointly required during parasite egress. CDPK1 and PKG play dominant roles in their respective modules, with CDPK2A and CDPK3 contributing less-essential functions. These supportive activities may be nonetheless important for parasite fitness under particular conditions. Additionally, these functional modules seem to be spatially distinct, with CDPK1/CDPK2A signaling occurring in the parasite cytoplasm, while CDPK3/PKG signaling occurs at the parasite PM (68). Understanding the topology of signaling pathways underlying key events in the parasite life cycle can help identify compensatory changes and predict phenotypic plasticity as we contemplate targeting parasite kinases for anti-parasitic therapies.

## MATERIALS & METHODS

### Parasite and host cell culture

*T. gondii* parasites were grown in human foreskin fibroblasts (HFFs) maintained in DMEM (GIBCO) supplemented with 3% heat-inactivated newborn calf serum (Millipore Sigma), 2mM L-glutamine (Thermo Fisher Scientific), and 10 μg/mL gentamicin (Thermo Fisher Scientific). Where noted, DMEM supplemented with 10% heat-inactivated fetal bovine serum (FBS, Millipore Sigma), 2mM L-glutamine (Thermo Fisher Scientific), and 10 μg/mL gentamicin was used. HFFs and *T. gondii* lines were monitored regularly and maintained as mycoplasma-free.

### Parasite transfection

Parasites were passed through 3 µm filters, pelleted at 1000 × *g* for 10 min, washed, resuspended in Cytomix (10⍰mM KPO_4_, 120⍰mM KCl, 150⍰mM CaCl_2_, 5⍰mM MgCl_2_, 25⍰mM HEPES, 2⍰mM EDTA, 2⍰mM ATP, and 5⍰mM glutathione), and combined with DNA to a final volume of 400 μL. Electroporation used an ECM 830 Square Wave electroporator (BTX) in 4 mm cuvettes with the following settings: 1.7 kV, 2 pulses, 176 μs pulse length, and 100 ms interval.

### Strain generation

Oligos were ordered from IDT. Primers, plasmids, and parasite strains used or generated in this study can be found in **Supplementary Table 1**. Descriptions of strain generation and plasmid construction, or relevant accession numbers, are also provided in the table.

### cDNA generation

Total RNA was extracted from *RH* parasites using Trizol. cDNA was generated according to package instructions for SMARTer PCR cDNA synthesis kit (Clontech/TakaraBio).

### Genomic DNA extraction

Extracellular parasites were pelleted at 1000 × *g* for 10 min and resuspended in phosphate buffered saline supplemented with Proteinase K (10 μg/mL). Suspensions were incubated at 37 for 1 h, 50 for 2 h, 95 for 15 min to extract genomic DNA.

### Immunoblotting

Parasite pellets were lysed in xenopus buffer (50 mM KCl, 20 mM HEPES, 2 mM MgCl_2_, 0.1 mM EDTA pH 7.5) supplemented with 1% TritonX-100, HALT protease inhibitor cocktail (ThermoFisher), and 10 μg/mL DNaseI (Sigma Aldrich) at room temperature for 1 hour with rotation. Lysates were combined with Laemmli buffer (for final concentration 2% SDS, 20% glycerol, 60 mM Tris HCl pH 6.8, 0.01% bromophenol blue) and 2-mercaptoethanol (1% final concentration) and boiled 10 min. Samples were run on a 7.5% SDS-PAGE gel (BioRad), transferred onto a nitrocellulose membrane in transfer buffer (25 mM TrisHCl, 192 mM glycine, 0.1% SDS, 20% methanol). Blocking and all subsequent antibody incubations were performed in 5% milk in TBS-T (20 mM Tris, 138 mM NaCl, 0.1% Tween-20). Primary and secondary antibody incubations proceeded for 1 h rocking at room temperature, with three TBS-T washes between primary and secondary and between secondary and imaging. Imaging was performed using a LI-COR Odyssey. Primary antibodies used were mouse anti-Ty (76) and rabbit anti-HA (71-5500, Invitrogen) or rabbit anti-TgACT1(77). Secondary antibodies were anti-mouse-800CW (LI-COR) or anti-rabbit-680RD (LI-COR).

### Live cell imaging

Parasites were inoculated onto glass-bottom 35mm dishes (Mattek) containing HFFs. At 24 h post-infection, intracellular parasites were imaged with an Eclipse Ti microscope (Nikon) with a 60X objective using the NIS elements imaging software and a Zyla 4.2 sCMOS camera. FIJI software was used for image analysis and processing.

### Immunofluorescence assays

Parasites were inoculated onto coverslips containing HFFs. At 24 h post-infection, intracellular parasites were fixed with 4% formaldehyde and permeabilized with 0.05% saponin in PBS. Nuclei were stained with Hoechst 33258 (Santa Cruz) and coverslips were mounted in Prolong Diamond (Thermo Fisher). Ty was detected using a mouse monoclonal antibody (76). HA was detected using a rabbit monoclonal antibody (71-5500, Invitrogen). Primary antibodies were detected with anti-mouse Alexa-Fluor 488 and anti-rabbit Alexa-Fluor 594 secondary antibodies (Invitrogen). Images were acquired with an Eclipse Ti microscope (Nikon) with a 60X objective using the NIS elements imaging software and a Zyla 4.2 sCMOS camera. FIJI software was used for image analysis and processing.

### Egress assays

Egress was quantified in a plate-based manner as in (55). HFF monolayers in a clear bottomed 96-well plate were infected with parasites and allowed to incubate 24 h before exchanging media for FluoroBrite DMEM (ThermoFisher) supplemented with 10% FBS and applying pre-treatments according to experiment type. Three images were taken before zaprinast (final concentration 500 μM; MilliPore Sigma) or A23187 (final concentration 8 µM; MilliPore Sigma) and DAPI (final concentration 5⍰ng⍰/mL) were added, and imaging of DAPI-stained host cell nuclei continued for 9 additional minutes before 1% Triton X-100 was added to all wells to determine the total number of host cell nuclei. Imaging was performed at 37°C and 5% CO_2_ using a Biotek Cytation 3 imaging multimode reader with a 4X objective. % egress was calculated as [(nuclei at time_n_ - nuclei at time_1_)/(nuclei at time_final_ – nuclei at time_1_)] * 100, and egress efficiency was normalized to egress of vehicle-treated parasites (% vehicle). Results are the mean of three wells per condition and are representative of at least three independent experiments. MOI was determined by performing immunofluorescence on a parallel plate of parasite-infected HFFs. Briefly, infected monolayers were fixed and permeabilized with 100% methanol for 2 min. Parasites were stained with either rabbit anti-TgALD (78) or guinea pig anti-CDPK1 antibody (79) and anti-rabbit or anti-guinea pig Alexa-Fluor 594 secondary antibody, and nuclei were stained with Hoechst 33258 (Santa Cruz Biotechnology). Imaging was performed using a Biotek Cytation 3 imaging multimode reader with a 20X objective, and parasite vacuoles and host nuclei were manually counted.

For AS kinase egress assays, HFFs were infected with 5 ×⍰10^4^ parasites per well and treated with 3 µM 3-MB-PP1 (MilliPore Sigma) or equivalent dilution of DMSO for 20 min prior to analysis.

For AID egress assays, HFFs were infected with 1 ×⍰10^5^ parasites per well of TIR1 parental or CDPK-AID lines and treated with 500 µM IAA or equivalent dilution of PBS for 3 h prior to analysis.

For epistasis endpoint assays, HFFs were infected with 1 ×⍰10^5^ parasites per well of TIR1 parental or CDPK-AID lines. Pre-treatment consisted of 500 µM IAA or equivalent dilution of PBS for 3 h, followed by 3-MB-PP1 (series from 2.5 µM to 0.039 µM) or equivalent dilution of DMSO for 30 min prior to analysis, and wells were stimulated to egress with zaprinast (final concentration 500 μM) for 20 min; all incubations were performed at 37°C and 5% CO_2_. Images were collected pre-stimulation (time_pre_), 10 min post addition of zaprinast (final concentration 500 μM) and DAPI (time_stim_), and 1 min post addition of 1% TritonX-100 (time_final_). % egress was calculated as [(nuclei at time_stim_ - nuclei at time_pre_)/(nuclei at time_final_ – nuclei at time_pre_)] * 100 and normalized to wells that did not receive IAA or 3-MB-PP1 for each strain. EC_50_ was determined by non-linear regression analysis performed using the non-sigmoidal dose-response with variable slope function in GraphPad Prism, with top constrained to 100 and bottom constrained to 0; significance was calculated using an F test to determine if all data (±IAA) fit to a single curve.

### Invasion assays

Briefly, parasite vacuoles were mechanically-dissociated and filtered through 5 µm filters, pelleted, and resuspended in invasion media (HEPES-buffered DMEM without phenol red) supplemented with 1% FBS. HFF monolayers in clear-bottom 96 well plates were incubated with parasite suspensions for 10 min at 37⍰ to stimulate invasion after centrifuging the plates at 290 x *g* and room temperature for 5 min. Wells were fixed with 4% formaldehyde and followed by antibody staining to differentiate between extracellular and total parasites and to detect nuclei. Samples were imaged using a Biotek Cytation3 imaging multimode reader with a 20X objective and imaging in montage mode.

For AS kinase invasion assays, parasites were pre-treated with 3-MB-PP1 (0.33 µM final concentration) or an equivalent dilution of DMSO in invasion media for 20 min at 37⍰, then 1 × 10^5^ parasites were added to 3 wells of a clear-bottom 96-well plate containing HFFs. Following incubation and fixation, extracellular parasites were stained with mouse anti-SAG1 antibody (80) conjugated to Alexa594. All parasites were stained by permeabilizing with 0.25% TritonX-100 and staining with anti-SAG1 antibody conjugated to Alexa488, and nuclei were stained with DAPI. The number of invaded parasites per field of view was counted using a size-based macro and normalized to the number of host cells in the same field of view (intracellular Tg/HCN). The final invasion efficiency for each replicate was normalized to the invaded parasites per host cell nuclei of the DMSO-treated parasites (% vehicle).

For AID invasion assays, parasite lines were each passed to two flasks of HFFs. 24 h pre-analysis one flask was supplemented with vehicle (PBS) and the second flask was supplemented with IAA to a final concentration of 500 µM. Following parasite harvest, 2 × 10^5^ parasites were added to 3 wells of a clear-bottom 96-well plate containing HFFs. Following incubation and fixation, extracellular parasites were stained with mouse anti-SAG1 antibody (80). All parasites were stained by permeabilizing with 0.25% TritonX-100 and staining with rabbit anti-GAP45, generated as previously described (81) and kindly provided by R.D. Ethridge (University of Georgia, Athens). Cells were subsequently stained with anti-rabbit Alexa594 antibody (Invitrogen), anti-mouse Alexa488 antibody (Invitrogen), and Hoechst 33258 (Santa Cruz Biotechnology). Images were acquired using a Cytation 3 imager (BioTek), and analyzed using custom FIJI macros to count the number of parasites and host-cell nuclei (49), plotted as intracellular Tg/HCN.

### Plaque assays

CDPK-AID and TIR1 parental parasites were inoculated into 6-well plates or 15 cm dishes of HFFs maintained in DMEM supplemented with 10% FBS and incubated overnight before supplementing with IAA to a final concentration of 500 µM or with PBS. Where indicated, plates were also supplemented with 3-MB-PP1 to a final concentration of 40 nM or with DMSO. Plates were allowed to grow undisturbed for 8 days then washed with PBS and fixed for 10 min at room temperature with 100% ethanol. Staining was performed for 5 min at room temperature with crystal violet solution, followed by two washes with PBS, one wash with water, and drying overnight. Plaques were counted manually, and plaque areas measured using FIJI software.

### FACS analysis

For IAA-induced depletion experiments, intracellular parasites were treated with either 500 µM IAA or an equivalent dilution of PBS for 1, 3, or 24 h. Following treatment, parasites were mechanically-dissociated by passing through a 27-gauge needle, isolated by filtration, and analyzed by flow cytometry with a Miltenyi MACSQuant VYB and plots were prepared using FlowJo software.

### Microneme protein secretion assays

CDPK-GLuc lines were each used to inoculate two flasks of HFFs. 24 h pre-analysis one flask was supplemented with vehicle (PBS) and the second flask was supplemented with IAA to a final concentration of 500 µM. Parasite vacuoles were mechanically-disrupted and parasites filtered through 5 µm filters, pelleted, and resuspended in cold intracellular buffer with free Ca^2+^ clamped at 100nM (ICB; 137mM KCl, 5mM NaCl, 20mM HEPES, 10mM MgCl_2_). 1 × 10^6^ parasites were combined with ICB, ICB supplemented with 3% FBS, or ICB supplemented with zaprinast (500 µM) or an equivalent dilution of DMSO with or without 3% FBS into 3 wells of 96 well round-bottom plate and incubated for 5 or 30 min at 37⍰ to stimulate secretion. Excreted/secreted antigen (ESA)-containing supernatants were collected by centrifugation at 1200 x *g* for 8 min at 4⍰ to pellet parasites. Parasite lysates were prepared according to Pierce Gaussia Luciferase Glow Assay Kit (ThermoFisher) and plated in triplicate with wells containing lysate from 6.6 × 10^5^, 2.2 × 10^5^, and 7.4 × 10^4^ parasite-equivalents for each strain with or without IAA. ESAs and lysates were plated in white-bottom assay plates (PerkinElmer #6002290) and incubated at room temperature for 5 min with substrate-containing assay solution before detecting luciferase signal using a Biotek Cytation3 multimode reader. For each independent replicate, background luminescence (from wells without parasites) was subtracted from ESA luciferase values.

### 3D Motility

Parasites in HFF cells were treated with either 500 µM auxin or PBS for three hours prior to harvest by mechanical dissociation. Pitta imaging chambers containing Hoechst 33342-stained parasites embedded in polymerized Matrigel were prepared as previously described (62). A Nikon Eclipse TE300 epifluorescence microscope (Nikon Instruments, Melville, NY) equipped with a 20× (0.65 pixel/µm) PlanApo λ (NA 0.75) objective and NanoScanZ piezo Z stage insert (Prior Scientific, Rockland, MA) was used to image the fluorescently-labeled parasite nuclei. Time-lapse video stacks were collected with an iXON Life 888 EMCCD camera (Andor Technology, Belfast, Ireland) using NIS Elements software v.5.11 (Nikon Instruments, Melville, NY) and pE-4000 LED illumination (CoolLED, Andover England). Images (1024 pixel × 384 pixel) were collected 1 µm apart in *z*, spanning 40 µm. Each *z* slice was imaged twice, at excitation wavelengths 385 nm (Hoechst) then 490 nm (mNeonGreen), each for 15 ms, before moving to the next *z* slice. The same volume was successively imaged 60 times at both wavelengths over the course of 78 seconds. The volume of each video stack was therefore 665.6 µm × 249.6 µm × 40 µm (*x, y, z*), and each dataset contained 2 × 60 stacks. The camera was set to trigger mode, no binning, readout speed of 30 MHz, conversion gain of 3.8x, and EM gain setting of 300.

Datasets were analyzed in Imaris ×64 v. 9.2.0 (Bitplane AG, Zurich, Switzerland). Parasites were tracked using the ImarisTrack module within a 1018 pixel × 380 pixel region of interest to prevent artifacts from tracking objects near the border. Spot detection used estimated spot diameters of 4.0 × 4.0 × 8.0 µm (*x, y, z)* for the fluorescently-labeled nuclei. A maximum distance of 6.0 µm and maximum gap size of 2 were applied to the tracking algorithm. Tracks with durations under ten seconds or displacements of less than 2 µm were discarded to avoid tracking artifacts and parasites moving by Brownian motion, respectively (62). Accurate tracking was confirmed by visual inspection of parasite movements superimposed on their calculated trajectories. The mNeonGreen signal was used to confirm the expected level of the mNeonGreen-Ty-AID-tagged CDPK protein in tracked parasites; parasites that were treated with auxin and remained positive for mNeonGreen were excluded from the analysis. Each experiment analyzed three biological replicates, each consisting of three technical replicates.

## Data Availability

Primers, plasmids, and parasite strains used or generated in this study can be found in **Supplementary Table 1**. Oligos, plasmids, and strains generated within this study are available from the corresponding author by request.

## ACKNOWLEDGEMENTS

We thank the L.D. Sibley (Washington University in St. Louis) for the TIR1 strain; S.M. Sidik for generation of the Cas9 nickase plasmid; B.M. Markus for generation of the mNeonGreen-mAID-Ty plasmid; R.D. Etheridge (University of Georgia, Athens) for the GAP45 antibody; VEuPathDB and all contributors to this resource. This work was supported by grants from the National Institutes of Health to S.L. (R01AI144369) and G.E.W. (R01AI139201), and training grant and fellowship support for R.V.S. (T32AI055402 and F31AI145214).

## AUTHOR CONTRIBUTIONS

E.S. and S.L. designed the overall study and experiments. C.G.H. generated and validated the AS-kinase strains and performed initial construction of complementing constructs. R.V.S. performed and quantified the gliding motility experiments. E.S. performed all remaining parasite strain construction and experiments. E.S. and S.L. wrote the manuscript and all authors reviewed, offered input, and approved the final draft.

